# Robot-based 3D-multispectral monitoring of soybean in a spatially heterogenous agrivoltaic environment

**DOI:** 10.64898/2026.03.31.715529

**Authors:** Avinash Agarwal, Christoph Jedmowski, Ilgaz Askin, Erekle Chakhvashvili, Matthias Meier-Grüll, Joschka Neumann, Michael Quarten, Uwe Rascher, Angelina Steier, Onno Muller

## Abstract

Agrophotovoltaic (APV) systems provide a unique opportunity for improving agricultural land-use efficiency by combining solar energy capture via photovoltaic panels with crop production. However, in-depth information on plant growth patterns within the spatially heterogenous microclimate created by the intermittent shading of APVs is largely missing. In the present study, we implement a customized robot-mounted 3D-multispectral imaging system to closely monitor the growth and spectral reflectance patterns of a conventional soybean cultivar “Eiko” (EK) and a chlorophyll-deficient mutant variety MinnGold (MG) under an APV system. Weekly trends in canopy morphometric features revealed significant variations in plant height, 3D leaf area, light penetration, and canopy volume across the APV field depending on the proximity with the overhead solar panels for both EK and MG, with plants receiving adequate rainfall and intermittent shade performing the best. Furthermore, although spectral indices exhibited variations between EK and MG due to intrinsic differences in pigmentation, symptoms of stress could be detected for both genotypes within rain-shaded areas of the APV plot. Hence, the present investigation depicts the potential for complementary usage of robotics and machine vision for high-precision high-throughput crop monitoring under APVs, which would enable better crop management within such non-homogenous cultivation systems.

## Introduction

The aim of Agrophotovoltaics (APVs), also known as agrivoltaics, is to amalgamate crop production and renewable energy generation by installing photovoltaic panels (PVPs) for solar energy capture on agricultural land, while maintaining adequate ground-clearance for regular agricultural activities underneath (Weselek et al., 2019). The aim of such systems is to boost energy security while maintaining food security, thereby improving land-use efficiency sustainably. Although the concept was introduced more than four decades ago by Goetzberger and Zastrow (1982), practical implementation has gained momentum in the past decade due to rising necessity for renewable energy and growing awareness of this unconventional farming practice (Weselek et al., 2019; Cosme and Vázquez y Parraguirre, 2025; Campana et al., 2025; Priya et al., 2026).

Despite the simplicity of the concept, integration of PVPs within contemporary agricultural frameworks necessitated the redressal of a major bottleneck. Since both plants and PVPs require solar energy for ensuring adequate productivity, the biggest hurdle was to resolve the competition for sunlight. To address this issue, pioneering studies within this field focused on two main aspects: 1) optimization of operational aspects of PVPs such as frequency, alignment, and ground-clearance within the field for adequate energy capture while ensuring sufficient sunlight reaches the crops, and 2) assessing the compatibility of such systems for diverse crops (Soto-Gómez, 2024; Mehta et al., 2025; Aly, 2026). Findings of such studies helped develop a better understanding related to feasibility and viability of APVs.

Till date, APV trials have already been conducted to assess the impact of overhead PVPs on the total yield using various crops, including rice (Lee et al., 2022; Nasukawa et al., 2025), maize (Amaducci et al., 2018; Ramos-Fuentes et al., 2023), soybean (Lee et al., 2022; Jo et al., 2022), tomato (AL-agele et al., 2021; Mohammedi et al., 2023; Savalle--Gloire et al., 2025), sweet potato (Maruyama et al., 2026), and sugarcane (Ferreira Junior et al., 2025). Many such studies reported a clear deviation in crop productivity under the PVPs as compared to the control plots, either positive or negative, with the main driver being altered crop microclimate within the APV plots (Priya et al., 2026). While change in light availability for plants under the PVPs is to be expected (Jedmowski et al., 2024; Magarelli et al., 2025; Priya et al., 2025), it was observed that these large overhead structures also affected climatic parameters such as air and soil temperature, relative humidity, soil moisture, airflow, and distribution of precipitation (Cosme and Vázquez y Parraguirre, 2025; Ferreira Junior et al., 2025; Zou et al., 2025). Since PVPs cause this “interference” physically, the impact on plant microclimate within such a system could be considered as a function of the position of each plant relative to the PVP installations in its vicinity.

Gaining knowledge of heterogeneity in plant growth environment is the next step for further optimizing APV operations, as it would allow better management of the system by focusing on factors such as precision agriculture, field planting design, mixed cropping, and modifications to the PVP layouts (Mehta et al., 2025). Although meticulous documentation of spatial variability in microclimate within APV systems could provide some hints towards potential plant growth responses (Magarelli et al., 2025; Priya et al., 2025; Zou et al., 2025), the results would be highly probabilistic for different crops and their diverse cultivars, ultimately requiring field tests for confirmation via labor-intensive manual measurements. Thus, a more straightforward solution would be to implement close-range high-throughput monitoring within such systems focusing on crops of interest. In this context, although machine vision technologies such as multispectral and 3D imaging have become increasingly popular in recent years for crop monitoring (Karukayil et al., 2026; Omia et al., 2026), their application for APV systems remains largely unexplored.

While a vast majority of studies related to APVs report final yield (Dinesh and Pearce, 2016; Jo et al., 2022; Lee et al., 2022; Potenza et al., 2022), which does not require in situ monitoring of crop traits, many of the studies that did report morphological and physiological characteristics, such as plant height, leaf dimensions, chlorophyll content, and photosynthetic activity (Weselek et al., 2021; Lee et al., 2022; Potenza et al., 2022; Ramos-Fuentes et al., 2023; Savalle--Gloire et al., 2025), provided the information based on manual or destructive measurements, which are labor intensive and have low throughput. Hence, deploying machine vision-based technologies via field-compatible robots within APV systems could boost research in this domain by enabling real-time high-throughput crop monitoring. This approach provides a potential link between detailed physiological investigations and large-scale remote sensing of crop performance by resolving crop responses at intermediate spatial scales.

To address this topic, we used a custom-made field-compatible robot mounted with a 3D-multispectral scanner to take a closer look at plant growth and spectral reflectance responses of two physiologically and structurally distinct soybean (*Glycine max*) genotypes, viz., “Eiko” (EK) which is a standard commercial soybean variety, and “MinnGold” (MG), a short, yellow-leaved mutant variety mainly used for experimental investigations (Campbell et al., 2015). The two genotypes were included based on their physiological and structural contrast within the same species (Sakowska et al., 2018; Acebron et al., 2023), creating the potential for exploring genotype-environment interactions while maintaining comparability (Supplementary Fig. S1). Plants were grown in a low-density elevated APV system and data was collected at weekly intervals for different locations within the APV field to generate a detailed spatiotemporal record of the vegetative growth phase. The study aims to elucidate the localized variations in plant traits due to environmental heterogeneity within an APV, while also depicting the potential for implementing state-of-the-art technologies such as machine vision and robotics for high-throughput in situ crop monitoring and management within APV-based systems.

## Methods

### Experimental design

The study was conducted over one full cultivation cycle between May and October 2025 at the Bürgewald APV site, North-Rhine Westphalia, Germany (50°51’49.87” N, 6°31’48.01” E; elevation above sea level: 122.5 m), managed by the Plant Sciences group, Institute for Bio- and Geosciences Plant Sciences (IBG-2), Forschungszentrum Jülich, Germany. The site lies within the Lower Rhine Plain (Niederrheinische Bucht), a region characterized by fertile loess-derived agricultural soils and intensive arable farming systems under predominantly rainfed conditions.

The experimental plot comprised of ca. 400 m^2^ planting area with four elevated PVP rows, and a control plot adjacent to it (Fig. 1A). The V-shaped PVP rows (length: 11 m, width: 3.6 m each) were positioned at a height of approx. 5.5 m from the ground, with a lateral spacing of 8.4 m, providing ca. 30% ground coverage, with a net horizontal inclination of 0° (Meier-Grüll et al., 2024). Briefly, each PVP system comprised of twenty bifacial silicon-based PV modules (dimensions: 1×2 m^2^) to provide 25 kW_peak_, and was connected to a battery pack with 35 kWh capacity. The electricity thus generated was used for charging the robot and all other devices used during the experiment.

**Fig. 1.**
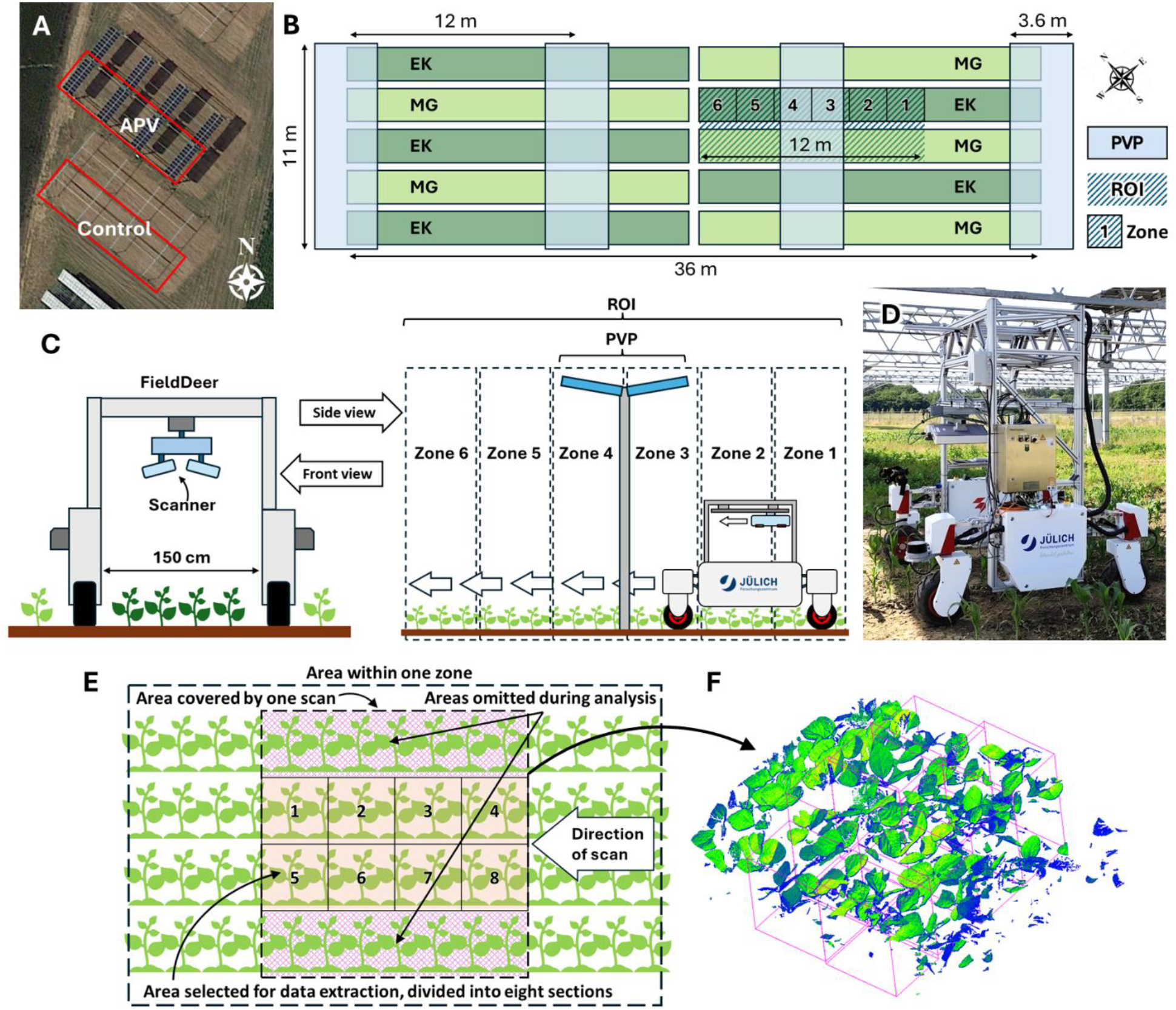
Satellite image of the Agrophotovoltaic (APV) and control plots (A), schematic overview of the experimental field depicting the planting layout along with the location of photovoltaic panels (PVPs) and region of interest (ROI; B), schematic representation of robotic operation for in-field data collection (C), the FieldDeer robotic system mounted with a 3D-multispectral scanner (D), schematic overview of scan preprocessing (E), and a representative 3D scan of soybean plants (F). Dark and light shades of green (B) indicate Eiko (EK) and MinnGold (MG) soybean genotypes, respectively.

Fertilization of soil and sowing of soybeans was carried out as per local guidelines, and plants were allowed to be irrigated by rainfall only. A weather station (Campbell Scientific, USA) installed at the experimental site allowed monitoring of climatic conditions. Average air temperature during the growth period (June–September 2025) was 18.2 °C, along with total precipitation 233 mm, and average soil temperature 18.1 °C. Spatial patterns in incoming solar radiation were recorded at hourly intervals using photosynthetically active radiation (PAR) sensors (LI-COR, USA) on a clear sunny day in the middle of the growth cycle (Supplementary Fig. S2A). Similarly, spatial variation in relative rainfall within the APV system was estimated by recording rainfall on 10 days using rain gauges placed at different locations relative to the PVPs (Supplementary Fig. S2B).

Seeds for both soybean genotypes were sown at a density of 30/m^2^ in the last week of May 2025, with the two varieties in alternate rows (Fig. 1B). A gap of ca. 50 cm was maintained between the 1.5 m wide rows for driving the robot (Fig. 1C). An identical planting scheme was followed for the control plot. Representative areas (length: 12 m, width: 1.5 m) were earmarked for each genotype as regions of interest (ROIs) within the APV plot such that one PVP module divided the ROI into two symmetrical parts (Fig. 1B, C). As an exploratory proof-of-concept study, we prioritized detailed monitoring of a representative area within the APV system rather than extensive spatial coverage across the entire field. To record spatial heterogeneity in plant traits within the APV system, each ROI was divided into zones 1 to 6 (L: 2 m, W: 1 m each) depending on the distance from the center of the PVP from South-East to North-West (Fig. 1C). Thus, zones 1 and 6 were furthest away from the PVP, zones 3 and 4 were situated directly under the PVP, and zones 2 and 5 were located at an intermediate distance from the PVP. For the control plot, data was collected weekly at fixed locations randomly selected within the non-bordering rows. Multiple zones were not chosen within the control considering homogenous environmental conditions in the absence of overhead structures.

### Robot design and 3D-multispectral scanning

A custom-made semi-autonomous robot (“FieldDeer”), based on the Thorvald platform (Saga Robotics, Norway), was developed to be operated within the experimental plots (Fig. 1C, D). The FieldDeer was equipped with remote control functionality along with four-wheel drive and four-wheel steering, enabling mobility and sufficient maneuvering capacity within the field. A minimalistic aluminum frame, minimizing shading to avoid triggering a shade response of plants, was constructed for the FieldDeer. The sensors and associated data processors along with a programmable remote-controlled horizontal conveyor system were attached to the frame. The robot was also mounted with an RTK-GPS system for precise geolocation as well as 3D LiDAR sensors for detecting obstructions and terrain mapping. In addition, the FieldDeer was operated using the open-source Robot Operating System (ROS), which allowed flexible programming of the platform. The conveyor system was mounted with a PlantEye F600 3D-multispectral scanner (Phenospex, The Netherlands) to function like a line-scanner (Agarwal et al., 2024). The conveyor was connected to a pair of ON/OFF switches that would activate the scanner over the desired horizontal distance of operation, and subsequently return the scanner to the starting position. Power was supplied by an integrated 48 V, 70 Ah lithium battery for operating the robot, the conveyor, the scanner, and the computer connected to the scanner.

Based on the field design, the height of the conveyor was adjusted to sweep an area of ca. 1 m^2^ per scan, following a semi-discontinuous “stop-scan-go” approach (Fig. 1C). Herein, the FieldDeer was driven to the desired plot, stopped over each zone within the ROI such that the scanner was oriented directly above the central row of soybean plants. Subsequently, the conveyor-scanner system was triggered remotely to scan the canopy while the robot remained stationary. The scanner simultaneously collected 3D spatial information using a laser sensor (λ_m_ = 940 nm, spectral half-width [SHW] = 1.5 nm), and spectral data for blue (B; λ_m_ = 475 nm, SHW = 20 nm), green (G; λ_m_ = 535 nm, SHW = 80 nm), red (R; λ_m_ = 629 nm, SHW = 20 nm), and near-infrared (NIR; λ_m_ = 735 nm, SHW = 20 nm) channels with the respective spectral sensors. The setup provided an approximate resolution of 0.7 mm X-axis, 1 mm Y-axis, and 0.2 mm Z-axis. The scanning system was equipped with in-built LEDs corresponding to each spectral waveband to illuminate the samples uniformly during scanning. After the scan was complete, the robot was driven to the next zone to repeat the process. Scans were performed weekly at the same spots from 8 July 2025 to 26 August 2025 to record temporal changes over eight weeks of vegetative growth.

### Data pre-processing and feature extraction

Scans were immediately transferred to a computer connected to the scanner and subsequently processed using the Hortcontrol software (Phenospex). The software performed the following actions: 1) color balancing based on internal calibrations; 2) background removal using height and color thresholds; 3) storage of scans as point-cloud (.ply) files (Agarwal et al., 2025). For each scan, covering an area of ca. 1 m × 1 m, a region of 25 cm on the left and right edges was omitted during data pre-processing to avoid border effects (Fig. 1E). The remaining area of the scan was divided into eight equal sections for extracting the morphometric and spectral features (Fig. 1E, F). The following morphometric features were obtained from the 3D point cloud (Table 1): canopy height, canopy 3D leaf area, projected canopy volume, light penetration depth ratio. Similarly, spectral features were obtained from the multispectral data as follows (Table 1): Green Leaf Index (GLI), Normalized Difference Vegetation Index (NDVI), Normalized Pigment Chlorophyll ratio Index (NPCI), and Plant Senescence Reflectance Index (PSRI).

**Table 1.**
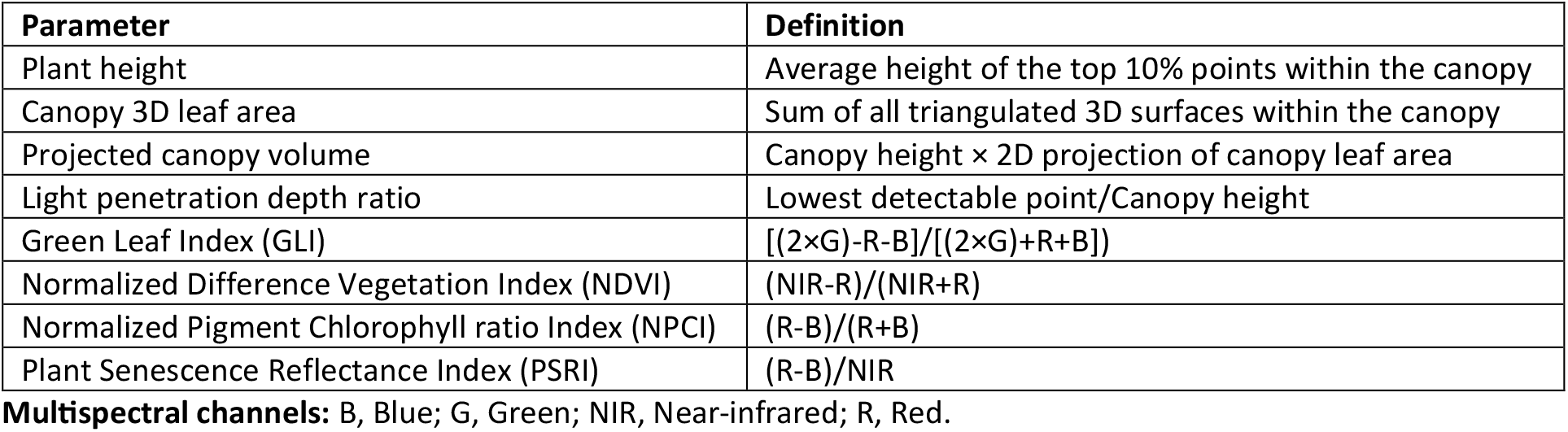
Morphometric and spectral features obtained from the 3D-multispectral scans.

### Statistical analysis

Weekly trends for the morphometric and spectral attributes of both genotypes were assessed for statistical significance in the R program (www.r-project.org) by using Robust Linear Mixed-effects Modeling (library: *robustlmm*) followed by pairwise Tukey’s post-hoc test. Herein, the “Robust” model was chosen as it is less sensitive to outliers. A quadratic temporal factor was incorporated to account for non-linear trends over the weeks. To obtain an overview of the entire period of observation, the model was designed to provide a marginal treatment effect over time. Sample identity was included as a random effect with both intercept and slope for time to account for repeated measurements. A simplified formula for the model is as follows:

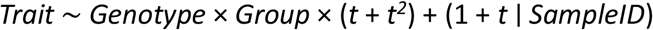

Here, *Trait* represents a unique plant attribute, *Genotype* indicates either EK or MG, *Group* denotes one of the APV zones or control, *t* denotes the temporal factor, and *SampleID* indicates a unique identifier assigned to each sampling section within a scan. Since the main aim of the study was to highlight the spatiotemporal heterogeneity in plant responses within APVs, “week” and “APV zone” were considered as the temporal and spatial factors, respectively, and data for both genotypes was handled independently. In addition to the *p*-values, which indicated the significance of difference, the analysis also yielded *z*-values representing the magnitude and direction (positive or negative) of deviation in the marginal trends. For the sake of simplicity, only the *p*- and *z*-values of the six zones compared to the control were used during visualization. Out of the 896 observations, i.e., 2 genotypes × 7 groups (control + six APV zones) × 8 weeks × 8 sample sections, 25 observations were omitted prior to analyses as erroneous measurements considering unusually high or low values for at least two attributes, clearly caused by technical interferences or inadequate exposure during scanning.

## Results

### Morphometric variations in soybean plants under APV conditions

Plant growth and spectral reflectance showed clear variations between the different APV zones as well as with the control, as depicted in the 3D scans from the eight week of observation (Fig. 2). In general, steady plant growth was recorded for both EK and MG in the control as well as APV conditions till the fifth week of observation, as indicated by the canopy volume, plant height, and 3D leaf area estimated from the scans (Fig. 3A, C, Supplementary Fig. S3). Concomitantly, LPDR presented an asymptotic decline with the passage of time for most samples, representing the increase in canopy density with plant growth (Fig. 3B, D). Although plant height for both genotypes exhibited a gradual increase in subsequent weeks as well, canopy volume and 3D leaf area showed a saturating or slightly declining trend from six weeks onwards.

**Fig. 2.**
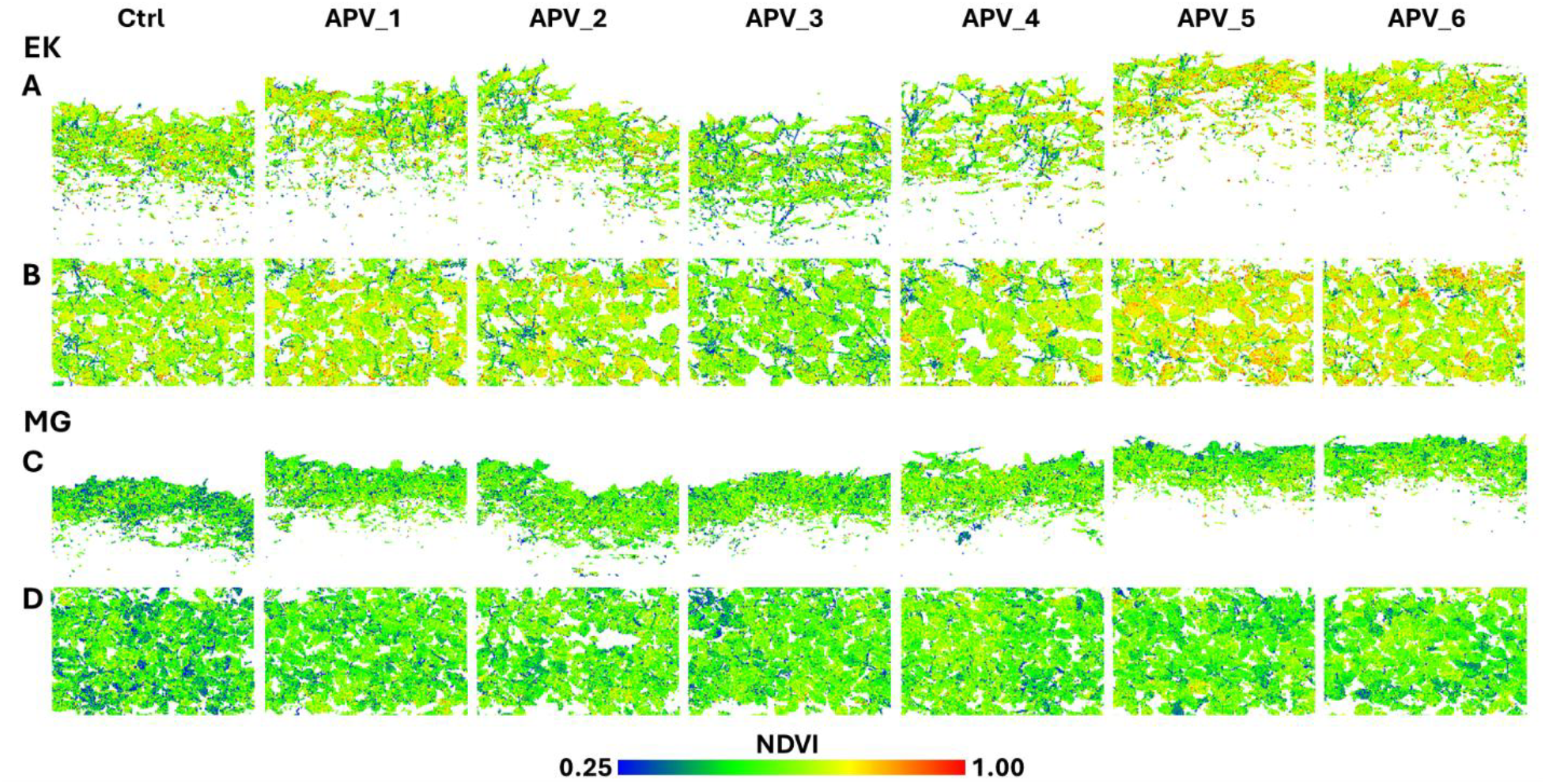
Side (A, C) and top (B, D) views for the 3D scans of Eiko (EK; A, B) and MinnGold (MG; C, D) soybean plants during the eight week of observation under the photovoltaic panels. APV_1 to APV_6: different zones within the Agrophotovoltaic plot; Ctrl: control; NDVI, Normalized Difference Vegetation Index.

**Fig. 3.**
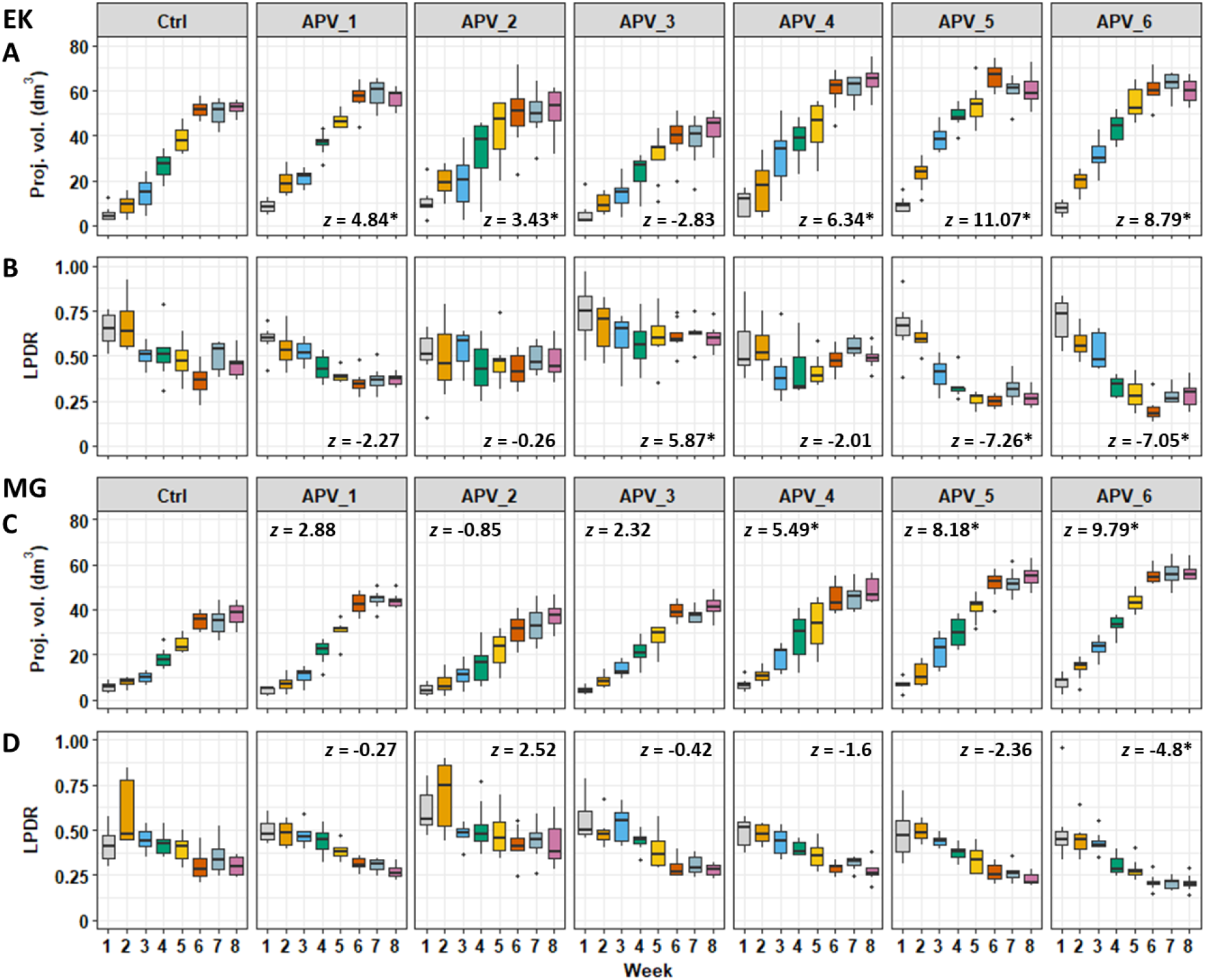
Weekly trends for morphometric attributes, viz., canopy projected volume (A, C) and light penetration depth ratio (LPDR; B, D), of soybean genotypes Eiko (EK) and MinnGold (MG) under photovoltaic panels. APV_1 to APV_6: different zones within the Agrophotovoltaic plot. Box-and-Whisker plots indicate the median (horizontal line), interquartile range (box), whiskers representing 5 and 95 % percentile, and the outliers (●) for *n* = 8 observations. Data indicated by * differed significantly (*p* < 0.05) from the control (Ctrl) as per Tukey’s test following Robust Linear Mixed-effects Modeling, with the *z*-values representing the magnitude and direction of deviation from the control.

For EK, canopy volume (Fig. 3A) and plant height (Supplementary Fig. S3B) indicated significantly better growth in APV zones 1, 2, 4, 5, and 6 as compared to the control (*p* < 0.05), with the strongest responses in zones 5 and 6 (*z* > 8). In contrast, both parameters suggested slightly poorer growth in zone 3 relative to control (*z* < -1.5; *p* > 0.05). Canopy 3D leaf area also indicated significantly better plant growth in zones 5 and 6 with respect to control (*p* < 0.05), although the differences were not significant for the other zones (Supplementary Fig. S3A). This trend was also reflected by the strong decline in LPDR for zones 5 and 6 (Fig. 3B), implying that the lower leaves became increasingly obscured over time as the canopy became more densely packed. Conversely, LPDR was significantly higher in zone 3 as compared to control (*z* = 5.87; *p* < 0.05) because of a sparser canopy owing to subdued growth.

Similar to EK, analysis of canopy volume, plant height, and 3D leaf area for MG also indicated superlative growth in zones 5 and 6 (*z* > 6.85; *p* < 0.05), followed by zone 4 (Fig. 3C, Supplementary Fig. S3C, D). However, unlike EK, plant height and canopy volume for zones 1 and 3 were only moderately higher than control, and slightly lower in zone 2 (*p* > 0.05). Suppression of growth for MG in zone 2 was also reflected by the relatively lower 3D leaf area compared to control (*z* = - 1.43; *p* > 0.05), whereas zones 1 and 3 had significantly higher, but intermediate values (*z* ∼ 3.2– 3.3; *p* < 0.05). The trend was corroborated by the LPDR of MG samples (Fig. 3D), wherein zone 6 exhibited significantly lower values than control (*z* = -4.8; *p* < 0.05) denoting a distinct increase in canopy compactness, followed by zone 5 (*z* = -2.36) and zone 4 (*z* = -1.59), whereas LPDR remained relatively higher within zone 2 (*z* = 2.52).

### Spectral variations in soybean plants under APV conditions

The numerical ranges and temporal trends for spectral indices varied considerably between EK and MG (Fig. 4, Supplementary Fig. S4) owing to the strong distinction in inherent leaf pigmentation (Supplementary Fig. S1). Hence, while GLI presented broadly homologous temporal trends with different ranges for both genotypes, the other three indices exhibited distinct variations over time between EK and MG. Specifically, GLI was generally found to increase from week 1 to week 5 for EK and till week 6 for MG, followed by a gradual decline across all the APV zones as well as the control (Fig. 4A, C). In contrast, NDVI peaked for EK around weeks 5 and 6, but continued increasing till week 8 for MG (Supplementary Fig. S4A, C), whereas NPCI showed a generally declining trend for EK, but peaked around weeks 4 and 5 for MG (Supplementary Fig. S4B, D). Moreover, although PSRI was found to reduce from week 1 to week 8 for both EK and MG, the decline was more prominent from week 1 to week 4 for EK, whereas it was more noticeable from week 5 to week 8 for MG (Fig. 4B, D).

**Fig. 4.**
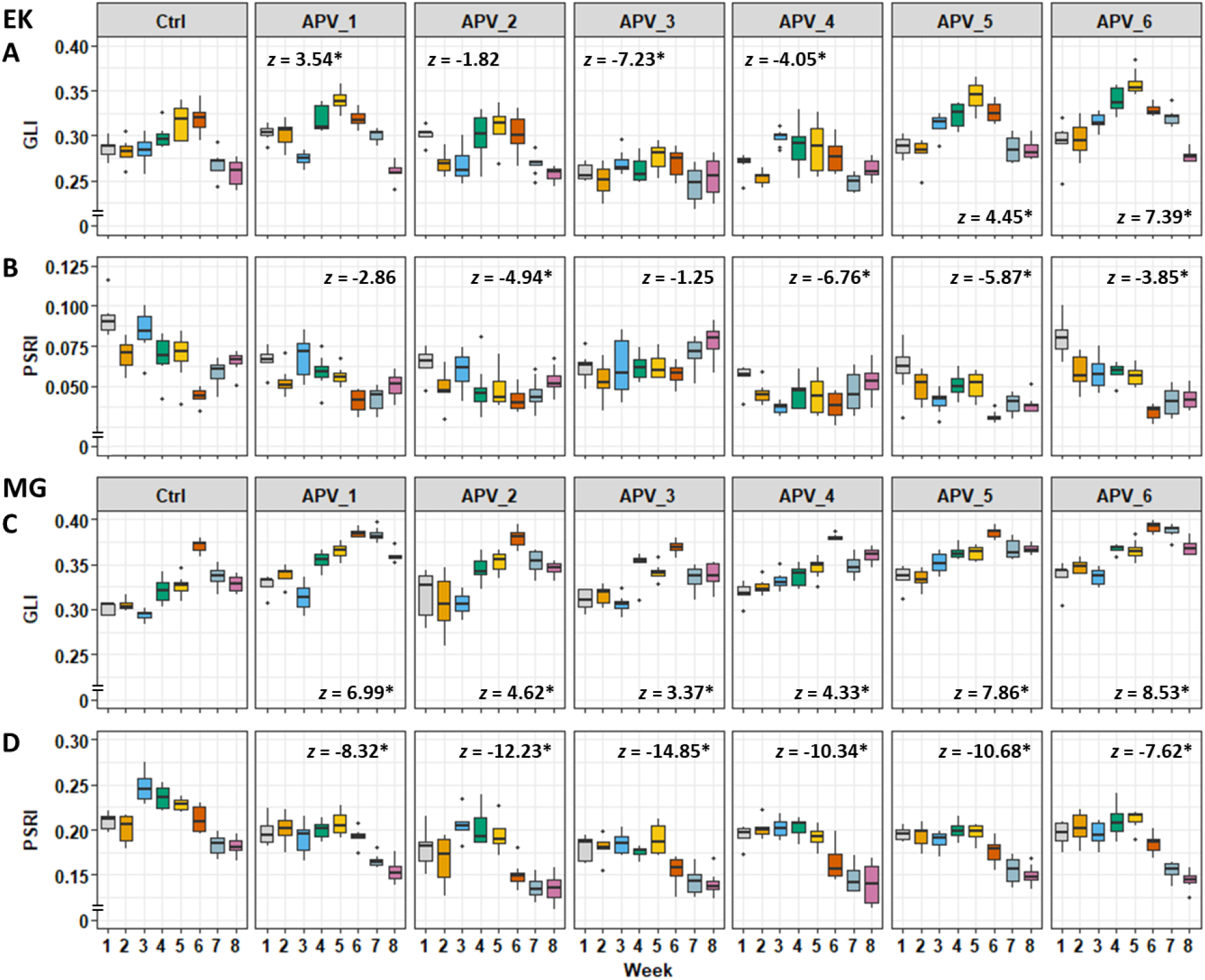
Weekly trends for spectral attributes, viz., Green Leaf Index (GLI; A, C) and Plant Senescence Reflectance Index (PSRI; B, D), of soybean genotypes Eiko (EK) and MinnGold (MG) under photovoltaic panels. APV_1 to APV_6: different zones within the Agrophotovoltaic plot. Box-and-Whisker plots indicate the median (horizontal line), interquartile range (box), whiskers representing 5 and 95 % percentile, and the outliers (●) for *n* = 8 observations. Data indicated by * differed significantly (*p* < 0.05) from the control (Ctrl) as per Tukey’s test following Robust Linear Mixed-effects Modeling, with the *z*-values representing the magnitude and direction of deviation from the control.

Amongst the different APV zones for EK, GLI showed the highest increment compared to control in zone 6 (*z* = 7.39, *p* < 0.05), followed by zones 5 and 1 (3.5 < *z* < 4.5, *p* < 0.05; Fig. 4A). While zone 2 exhibited a modest reduction in GLI with respect to control (*z* = -1.82), the index was significantly lower for zone 4 (*z* = -4.05, *p* < 0.05), and lowest for zone 3 (*z* = -7.23, *p* < 0.05). Similar trends were observed for NDVI in EK, wherein the index increased most distinctly for zones 5 and 6 (*z* > 8, *p* < 0.05), but remained significantly low compared to control in zone 3 (*z* = -3.35, *p* < 0.05; Supplementary Fig. S4A). NDVI was also significantly higher than control for EK in zones 1, 2, and 4 (4.5 < *z* < 5.5, *p* < 0.05), but remained lower than zones 5 and 6. Unlike GLI and NDVI, NPCI values for the different APV zones of EK were consistently lower than the control, although the spatial trends were highly diverse (Supplementary Fig. S4B). Specifically, NPCI was most significantly reduced in zone 4 (*z* = -8.93, *p* < 0.05), followed by zones 2, 5, and 3 (-6.4 < *z* < -4.5, *p* < 0.05), but did not differ significantly from control for zones 1 and 6. PSRI was also lower than control for all APV zones of EK (Fig. 4B), with the values being lowest in zone 4 (*z* = -6.76, *p* < 0.05), followed by zone 5 (*z* = -5.87, *p* < 0.05) and zone 2 (*z* = -4.94, *p* < 0.05).

Like EK, GLI of MG samples had the strongest increment in zone 6 (*z* = 8.53, *p* < 0.05), zone 5 (*z* = 7.86, *p* < 0.05), and zone 1 (*z* = 6.99, *p* < 0.05) with respect to control (Fig. 4C), whereas the values were moderately higher than control in zones 2, 3, and 4 (3.3 < *z* < 4.7, *p* < 0.05). In contrast, NDVI of MG samples for all APV zones was found to be considerably higher than the control (*z* > 6.5, *p* < 0.05), but did not differ significantly amongst the different zones (*p* > 0.25; Supplementary Fig. S4C). While NPCI for all APV zones of MG was lower than the control (Supplementary Fig. S4D), zone 3 recorded the lowest values (*z* = -11.37, *p* < 0.05), followed by zones 2 and 4 (*z* < -7, *p* < 0.05). Zones 1 and 5 also showed significantly lower NPCI (*z* < -3, *p* < 0.05), but did not differ significantly from the control in zone 6. In contrast, PSRI of MG was significantly lower than control in all APV zones (Fig. 4D), with the strongest response in zone 3 (*z* = -14.85, *p* < 0.05), followed by zones 2, 4, and 5 (*z* ∼ 10.3–12.5, *p* < 0.05).

## Discussion

### Impact of APV environmental heterogeneity on plant traits

Assessment of morphometric and spectral attributes of EK and MG soybean genotypes (Figs. 3, 4, Supplementary Figs. S3, S4) clearly highlighted the diversity in plant responses across different locations within the APV plot. Notably, plant growth was found to be consistently better than the control plot within APV zones 5 and 6 for both genotypes (Fig. 3, Supplementary Fig. S3), whereas EK samples in zone 3 and MG samples in zone 2 showed perceptible repression of growth. Because plants in both the control and APV plots were grown under comparable management conditions, the observed spatial differences in growth were likely associated with localized microclimatic variations induced by the PVPs.

On-field PVP installations have been reported to affect various climatic factors that impact plant growth (Ferreira Junior et al., 2025; Zou et al., 2025; Magarelli et al., 2025; Priya et al., 2025). Hence, to better understand the reason behind the heterogeneity in growth responses within our study, various climatic parameters were monitored as well. Diurnal trends in PAR recorded at hourly intervals for the different APV zones revealed that although the total amount of sunlight reaching the APV plots throughout the day was marginally lower than the control plots (Supplementary Fig. S2A), the mean diurnal PAR was consistent across the different APV zones. Moreover, spot measurements of air temperature and relative humidity presented only minor differences between the APV field and the control plot (data not shown). This is understandable considering the minimal restriction on air movement due to the high ground clearance (∼5.5 m) and low ground coverage (ca. 30%) of the present APV system.

In contrast, a clear difference was found in the amount of rainfall each APV zone received (Supplementary Fig. S2B). Specifically, APV zones 5 and 6 were found to receive rainfall at similar levels as the control area, whereas zone 3 faced the sharpest decline, followed by zone 2, both receiving ∼50% average relative rainfall compared to the control. While reduction in rainfall for zone 3 is understandable considering its location directly under the PVP, the decline in rainfall for zone 4, which also lies under the PVP, was not as strong. Instead, zone 2, which does not lie directly beneath the PVP, faced a sharper decline than zone 4. This was potentially due to persistent South-Western winds which shifted the rain-shadow region Eastwards from the expected area under zones 3 and 4 towards the area under zones 2 and 3. Since the experimental sites were exclusively irrigated by rainfall, spatial variations in precipitation within the APV field affected soil moisture content and water availability for the plants. Hence, the trends of relatively subdued growth for the two soybean genotypes in the respective zones of the APV field could be due to reduced water availability.

Even though plants had unimpeded rainfall in the control plot, plant growth responses were comparable to or lower than that observed in some APV zones. In contrast, plants within APV zones 5 and 6, which received comparable amounts of rainfall as the control along with intermittent shading from the PVPs, showed the best growth responses amongst all groups (Figs. 2, 3, Supplementary Fig. S3). Hence, transient shading within the APV system could be deemed advantageous for the soybeans, potentially due to better leaf water retention along with intermittent respite from light and heat stress, as observed in various other crops as well (Amaducci et al., 2018; Ferrara et al., 2023; Ramos-Fuentes et al., 2023).

Possibility of lower physiological stress for soybean within specific APV zones was further substantiated by GLI, NDVI, NPCI, and PSRI spectral index values (Fig. 4, Supplementary Fig. S4). It must be noted that while plant spectral indices represent relative changes in leaf reflectance across various wavebands, they provide an insight into the physiological status of the plants owing to their dependence on leaf pigmentation and healthy biomass content (Carter and Knapp, 2001; Berger et al., 2022; Kior et al., 2024). Accordingly, GLI and NDVI have been utilized as positive indicators of plant growth, pigment content, photosynthetic capacity, and health status, whereas NPCI is negatively correlated to chlorophyll content, and increasing PSRI has been associated with declining plant health (Peñuelas and Filella, 1998; Louhaichi et al., 2001; Merzlyak et al., 2002; Zarco-Tejada et al., 2005; Agarwal and Dutta Gupta, 2018; Tayade et al., 2022; Agarwal et al., 2025). Thus, higher GLI and NDVI in zones 5 and 6 along with lower NPCI and PSRI compared to control (Fig. 4, Supplementary Fig. S4) clearly indicated that plants within these areas of the APV system had higher photosynthetic pigment content and healthier biomass. Conversely, plants in the control plot and APV zones 2 and 3 with subdued growth exhibited signs of borderline stress, potentially due to overexposure to sunlight and reduced water availability, respectively.

Previous studies with different soybean varieties have reported reduction in growth and yield under APVs as compared to the control (Lee et al., 2022; Jo et al., 2022; Hu et al., 2024). However, Potenza et al. (2022) clearly depicted the spatial heterogeneity in soybean growth responses within the APV field, with certain APV areas showing comparable growth as the control samples. Furthermore, earlier investigations on soybean response to shading have indicated that moderate shading had a positive effect on plant growth (Bing-xiao et al., 2020) as well as photosynthetic potential (Yao et al., 2017). Additionally, the study by Zhang et al. (2016) specifically demonstrated that partial shading with adequate irrigation was beneficial for soybean, whereas shading with water stress was especially detrimental. This is in agreement with our observation of spatially contrasting growth responses between different APV zones due to heterogenous shade–rainfall interactions. Hence, zones receiving intermittent shading with adequate rainfall (zones 5 and 6) showed enhanced plant growth, whereas in shaded zones with reduced rainfall (zones 2 and 3) comparatively suppressed plant growth was observed.

### Compatibility of robotics and machine vision with APV systems

While the above discussion explicates spatial heterogeneity in soybean traits across the APV system and its potential causes, it also underlines how a robot-mounted plant imaging system can be particularly helpful in close-range crop monitoring under PVPs. Specifically, identification of zones with comparatively reduced plant growth through robot-based spatial phenotyping may support precision management approaches in APV systems, including targeted irrigation, genotype choice, and adaptive light management in tracking installations. Although robots and machine vision have been widely explored and implemented for open-field as well as controlled-environment cultivation (Fontani et al., 2025), the concept has not been extensively tested within APV systems.

The past decade has witnessed an exponential increase in APV research, with strong emphasis on the assessment of crop responses, energy capture, and socio-economic variables to establish practically feasible models (Ghasemi and Sadeghkhani, 2025). Moving beyond these foundational domains of APV research, the next step would be the implementation of latest technologies for further advancing APV operations (Abidin et al., 2022; Klokov et al., 2023). As depicted in the present study, use of robots and machine vision provides an ideal example of implementing state-of-the-art technology within APV systems for mapping plant response patterns and identifying zones of interest that would require mitigation. While such an approach provides better insights into the impact of PVPs in agricultural fields and creates a scope for optimal crop management, the biggest advantage of deploying such technologies within APV fields is the readily available source of energy anywhere within the field.

Normally, agricultural robots are powered by batteries and hence need to be charged prior to operation via conventional electrical installations (Fontani et al., 2025). This additional energy requirement creates a financial load on cultivators during large-scale operations. However, an APV system creates the ideal synergistic solution for implementing robotics in agriculture, as it practically nullifies the bottlenecks associated with energy requirements for field robotics. Specifically, the APV system acts as a perpetual source of energy for the robot, with the possibility of in-field recharging. Keeping in mind this complementarity of APVs and robotics, the FieldDeer was exclusively charged using the energy produced by the PVPs installed over the experimental fields during the entire investigation. Moreover, the peripheral accessories used in the experiment, viz., the 3D-multispectral scanner, remote control, and the computer associated with the robot, were also powered by the same battery. Hence, from the perspective of energy requirement, APVs could be deemed as a very favorable cultivation practice for using robots and other modern agricultural technologies.

### Limitations and future perspectives

By design, the current study had two main objectives: 1) assessing variations in crop traits across the APV field, and 2) testing the feasibility of operating a robot mounted with a 3D-multispectral scanner for close-range high-throughput crop monitoring under APVs. As indicated by our findings, deploying robots for crop monitoring under PVPs creates a new avenue for APV research by enabling detailed assessment of crop responses within such systems. Owing to the highly exploratory nature of the present investigation, a focused experimental plan was adopted to achieve these goals simultaneously.

Future trials with multiple identical ROIs spread across the APV field could help validate the findings related to spatial heterogeneity in plant traits within such a system more robustly. Further, use of smaller zones (ca. 1 m) could provide more precise details about spatial variations in plant growth, which would be beneficial for tailoring APV crop management more accurately. Additionally, more intensive monitoring of the microclimate within the APV plot via numerous fixed sensors for continuous data collection would help map the microclimatic heterogeneity more precisely, providing a solid base for interpreting plant responses more specifically. While we have refrained from discussing attributes such as yield and photosynthetic activity considering the scope of the present topic, these aspects could also be addressed in future reports.

Furthermore, the current trial was carried out with only one type of PVP, one planting density, and two genotypes of the same crop species. However, testing different PVP designs with multiple planting densities and diverse crop species having varying light and water requirements would further highlight the potential for robot-based crop monitoring within APV systems and will be important to investigate the generality of the observed spatial growth patterns. Use of autonomous robot navigation and unmanned data acquisition using additional sensors such as thermal and hyperspectral cameras could further enhance the capability of proximal phenotyping approaches. While this study focused on plot-level heterogeneity, the methodology presented here has the potential to contribute to phenotyping across spatial scales by linking localized crop responses with broader monitoring applications. Overall, our findings emphasize that spatially heterogeneous cultivation environments, such as APV systems, require high-resolution monitoring approaches to enable site-specific crop management.

## Acknowledgements

We would like to thank Marcel Weckbecker, Bente Königs, and Yannis Grosch for their support with technical and field activities.

## Author contributions

AA: Methodology, Investigation, Formal analysis, Data curation, Software, Visualization, Writing - Original draft; CJ: Methodology, Investigation, Writing - reviewing and editing; IA: Resources, Methodology, Software, Data curation; AS: Resources; MQ: Resources; EC: Investigation; JN: Resources; MMG: Supervision, Funding acquisition, Project administration; UR: Supervision, Funding acquisition, Project administration; OM: Conceptualization, Supervision, Funding acquisition, Project administration, Writing - reviewing and editing.

## Data availability

All data supporting the findings of this study are available within the paper and within its supplementary data published online.

